# Paradoxical Roles of Peritumoral Myofibroblasts in Intrahepatic Cholangiocarcinoma Growth and Metastasis

**DOI:** 10.1101/2021.02.19.432035

**Authors:** Cheng Tian, Liyuan Li, Yizhen Li, Li Fan, Anthony Brown, Michelle Morrison, Eric J. Norris, Jun J. Yang, Evan S. Glazer, Liqin Zhu

## Abstract

Intrahepatic cholangiocarcinoma (iCCA) is an aggressive type of primary liver cancer characterized by its highly desmoplastic stroma. Compared to the ample knowledge on cancer-associated fibroblast residing within iCCA tumor mass, little is known about the fibrotic response from the tumor-surrounding liver and its role in iCCA development. In this study, we find a significant accumulation of peritumoral myofibroblasts (pMFs) in both patient iCCA and tumors from an iCCA orthotopic allograft mouse model. Using tumor and liver spheroid coculture we show that iCCA-liver interaction induces rapid pMF accumulation at the interface. We find pMFs placed around iCCA spheroids exert a strong suppressive effect on tumor cells growth, in contrast to the pro-proliferative effect of MFs mixed within tumor spheroids. However, prolonged iCCA-pMF interaction elicits tumor cell dissemination in vitro. We find an upregulation of Vcam1 in tumor cells in the early phase of iCCA-pMF interaction both in vitro and in vivo which is downregulated when tumor cells disseminate. Blocking Vcam1 activity in iCCA allograft mouse models slows primary tumor growth but lead to increased tumor metastasis. Our data suggest that pMFs are beyond simple pro- or anti-tumorigenic in iCCA, with the ability to suppress tumor growth but elicit tumor cell dissemination.

## INTRODUCTION

Cholangiocarcinoma is the second most common hepatic tumor after hepatocellular carcinoma (HCC). It is believed to originate from the biliary tracts within or outside the liver which lead to the development of intrahepatic or extrahepatic cholangiocarcinoma (iCCA or eCCA, respectively). iCCA has a worse prognosis than eCCA and is one of the deadliest cancers overall (1, 2). Approximately one-third of iCCA patients present with metastases at the diagnosis and up to two-third of patients experience and eventually succumb to relapse. Modern therapy has been designed based on iCCA tumor biology but only marginally increased patient overall survival thus far (3, 4).

One of the most prominent characteristics of iCCA is its highly desmoplastic stroma. This has led to investigations on cancer-associated fibroblasts (CAFs) as well as hepatic stellate cells (HSCs), the major cell type giving rise to liver myofibroblasts (MFs) via their activation (5), in iCCA progression (6–10). These studies mainly focus on CAFs and activated HSCs (aHSCs) that reside within iCCA tumor mass, and have consistently found these fibrotic components promote iCCA progression. Interestingly, recent studies on HCC patients, although limited, revealed that aHSCs and MFs are also present in the tumor-surrounding liver and are positively associated with HCC metastasis and recurrence (11, 12). Indeed, the liver is one of the few internal organs that are developmental equipped with a complex damage response machinery in order to protect its vital function in metabolism and immunity (13). Nearly every chronic liver condition results in liver fibrosis eventually (14). Therefore, it is conceivable that a growing malignant mass will elicit fibrotic responses in the peritumoral liver during its progression, resulting in a dynamic interaction between tumor and peritumoral MFs (pMFs) that feeds back into tumorigenesis. Compared to HCC, pMFs have not been well studied in iCCA. We reason that investigating iCCA pMFs will serve as one of the first efforts to include host response into the development of this rare but highly aggressive cancer. To do so, we acquired a cohort of iCCA patient tumors resected without neoadjuvant treatment and examined the distribution of their MFs with a focus on those in the peritumoral liver. We then tracked the spatiotemporal MF activation in an orthotopic allograft model of metastatic iCCA we established in the laboratory. To assess the impact of pMFs on iCCA progression, we developed multiple novel coculture systems to model tumor-pMF interaction in vitro. Lastly, we investigated the potential underlying molecular mechanisms that mediate the dynamic iCCA-pMF interaction.

## RESULTS

### Accumulation of pMFs in iCCA patient and mouse allograft tumors

To examine MFs in iCCA patient tumors, we performed IHC of alpha-smooth muscle actin (αSMA), a well-recognized MF marker, on iCCA patient tumors. To avoid therapy-induced liver fibrosis, we selected three tumors resected with no neoadjuvant treatment and with >=5 mm tumor-surrounding liver attached. We defined peritumoral as regions within 100 μm from the tumor border and tumor core as >=2mm from the border. In these tumors, we found a large number of αSMA^+^ MFs in the tumor core as well as in the tumor-surrounding liver (**Figure 1A)**. In particular, there was a consistently higher density of pMFs at the tumor border in all three tumors compared to the tumor core.

**Figure 1.**
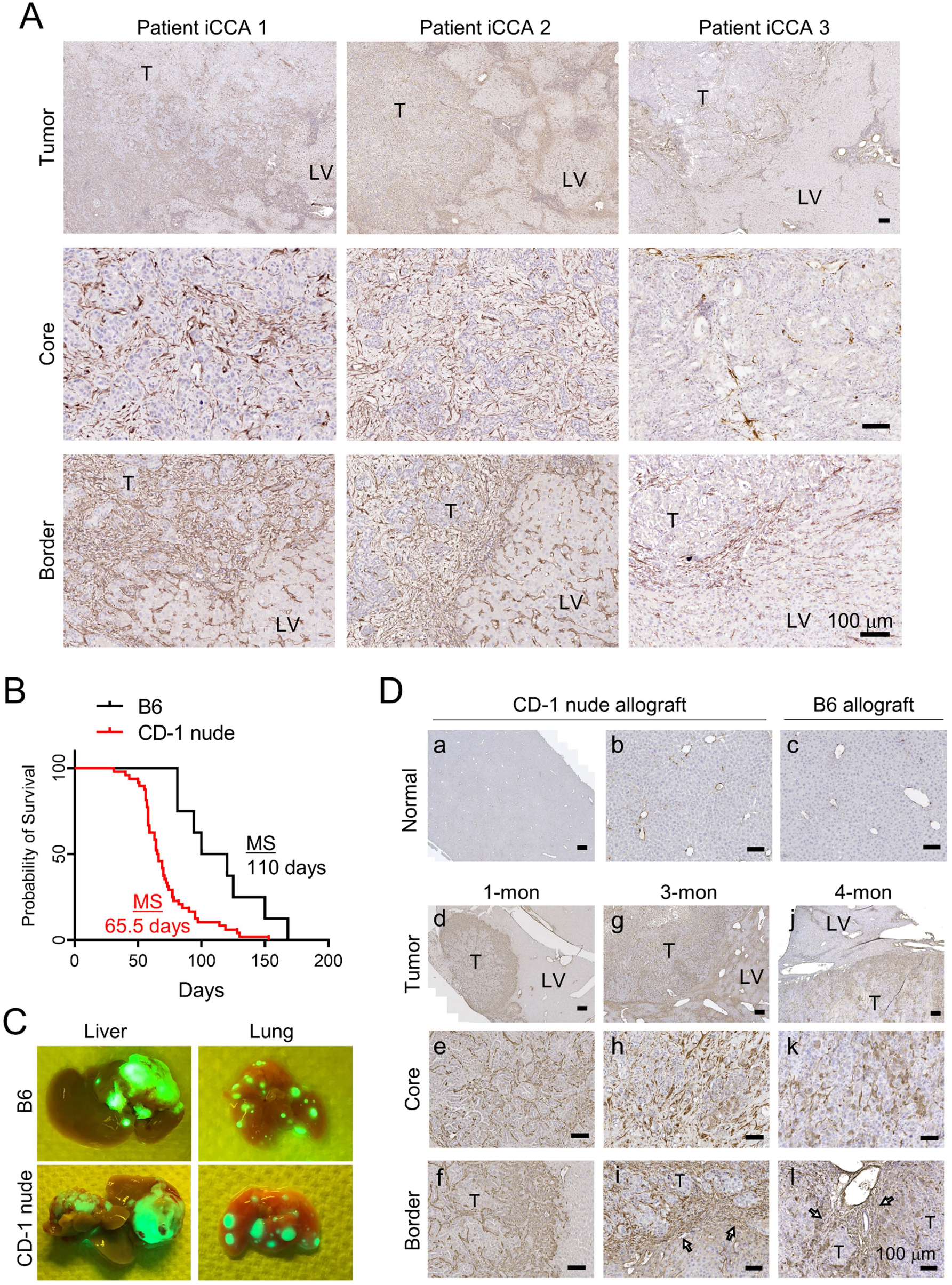
Accumulation of pMFs in mouse and patient iCCA tumors. **(A)** IHC of αSMA in three low-risk iCCA patient tumors showing higher density of MFs in the peritumoral region than the tumor core. **(B)** The animal survival curves of CD-1 nude and B6 mice orthotopic transplanted with PPTR tumor cells. **(C)** Gross GFP fluorescence images of the liver and lung from the iCCA allograft models showing the intrahepatic and lung metastases. **(D)** IHC of αSMA of the liver isolated from the indicated wildtype mice and allograft model at different time points. All scale bars are 100 μm.

Since MF activation in iCCA patients could be triggered either by tumorigenesis or by other non-tumor-related preexisting conditions, we utilized an orthotopic allograft model of metastatic iCCA we established in our laboratory to track iCCA-induce MF activation. Using tumor cells cultured from a previously reported *<u>P</u>rom1^CreERT2^; <u>P</u>ten^flx/flx^; <u>T</u>p53^flx/flx^; <u>R</u>osa-ZsGreen* (PPTR) liver cancer genetic model (15, 16), we developed an orthotopic allograft mouse models of metastatic iCCA in both CD-1 nude and B6 mice, with tumors developing faster and more consistently in the former (**Figure 1B** and **Suppl Figure S1A**). The *Rosa-ZsGreen (ZsG)* reporter allele in the PPTR tumor cells enabled direct visualization of gross tumor metastasis as well as disseminated tumor cells (DTCs) (**Figure 1C** and **Suppl Figure S1B**). We collected the wildtype livers and iCCA allograft tumors from different time points and performed αSMA IHC. As expected, αSMA positivity was only found in the smooth muscle cells lining large vessels in the normal mouse liver (**Figure 1D, a-c**). In the early-stage iCCA tumors collected after one month from CD-1 nude mice, we found a large number of αSMA^+^ pMFs at the tumor border similar to patient tumors (**Figure 1D, d**). Alpha-SMA^+^ MFs were also seen in the tumor core, however, at a significantly lower number than those in the peritumoral region (**Figure 1D, e & f**). In the 3-month tumors collected from the CD-1 nude allografts, there was a marked increase of αSMA^+^ MFs within the tumor core (intratumoral MFs, or iMFs) (**Figure 1D, g-h**). Accumulation of pMFs at the tumor border became more heterogeneous, however, was consistently found at the invasive border where dissociating tumor clusters were emerging (**Figure 1D, i**). Tumors collected at the end point from B6 allografts showed similar presence of MFs in both the tumor and surrounding liver, as well as pMF accumulation at the invasive border (**Figure 1D, j-l**). Interestingly, we also noticed widespread αSMA^+^ MFs in the distant liver far from the tumor mass in late-stage tumors, suggesting a systemic fibrotic reaction in the liver triggered by iCCA development. Overall, these observations in the iCCA orthotopic allograft model suggest a spatiotemporal MF activation in the host liver induced by iCCA development. Since the development of the orthotopic tumors was fasters and more consistent in CD-1 nude mice than B6 mice, all following in vivo experiments were performed in CD-1 nude mice.

### Tumor-liver spheroid coculture recapitulates pMF accumulation at the tumor border

Since tumor-associated changes in vivo could be part of a systemic response shaped by both the liver and other non-hepatic components, we developed an in vitro tumor-liver spheroid coculture model to detemine if the pMF accumulation we observed at the tumor border was induced directly by iCCA-liver interaction. Sperhoid models of liver hepatocytes (HCs) have been widely used to study HC metabolism and drug response (17–19). But reports on liver spheroid models consisting of both parenchymal and non-parenchymal compartments and their applications on cancer research are limited. We reasoned such models could be a powerful tool to dissect liver host response to tumorigenesis. To do so, whole liver cells (WLCs) were isolated from *Rosa^tdTomato/GFP^ (mTmG)* mice and cultured in AggreWell400 (AW400) plates for 7 days to generate spheroids (**Figure 2A**). Cells from mTmG mice expressed a strong *tdTomato* (TdT) red fluorescence protein (RFP). PPTR tumor cells were cultured in the same way and they formed homogeneous spheroids within two days (**Figure 2B**). IHC characterization of WLC spheroids confirmed the presence of HNF4α^+^ HCs, CD68^+^ Kupffer cells, CK19^+^ cholangiocytes and a small number of αSMA^+^ MFs which were likely given rise by HSCs activated by liver dissociation and culture (**Figure 2C**). To model tumor-liver interaction, WLC spheroids were mixed with tumor spheroids at a ratio of 15:1 and of WLC spheroid-only culture was used as the control (**Figure 2D**). All spheroids aggregated by 48 hours and the co-spheroids were maintained in culture until Day 7. The co-spherhoids were then collected and subjected to αSMA^+^ IHC. Compared to the small number and randomly distributed αSMA^+^ MFs in the WLC-only co-spheroids, WLC-tumor co-spheroids showed an evident accumulation of αSMA^+^ MFs at the interface of ZsG^+^ tumor spheroids and TdT^+^ WLC spheroids (**Figure 2E**), supporting the notion that pMF accumulation at iCCA tumor border in vivo was likely triggered by tumor-liver interaction directly.

**Figure 2.**
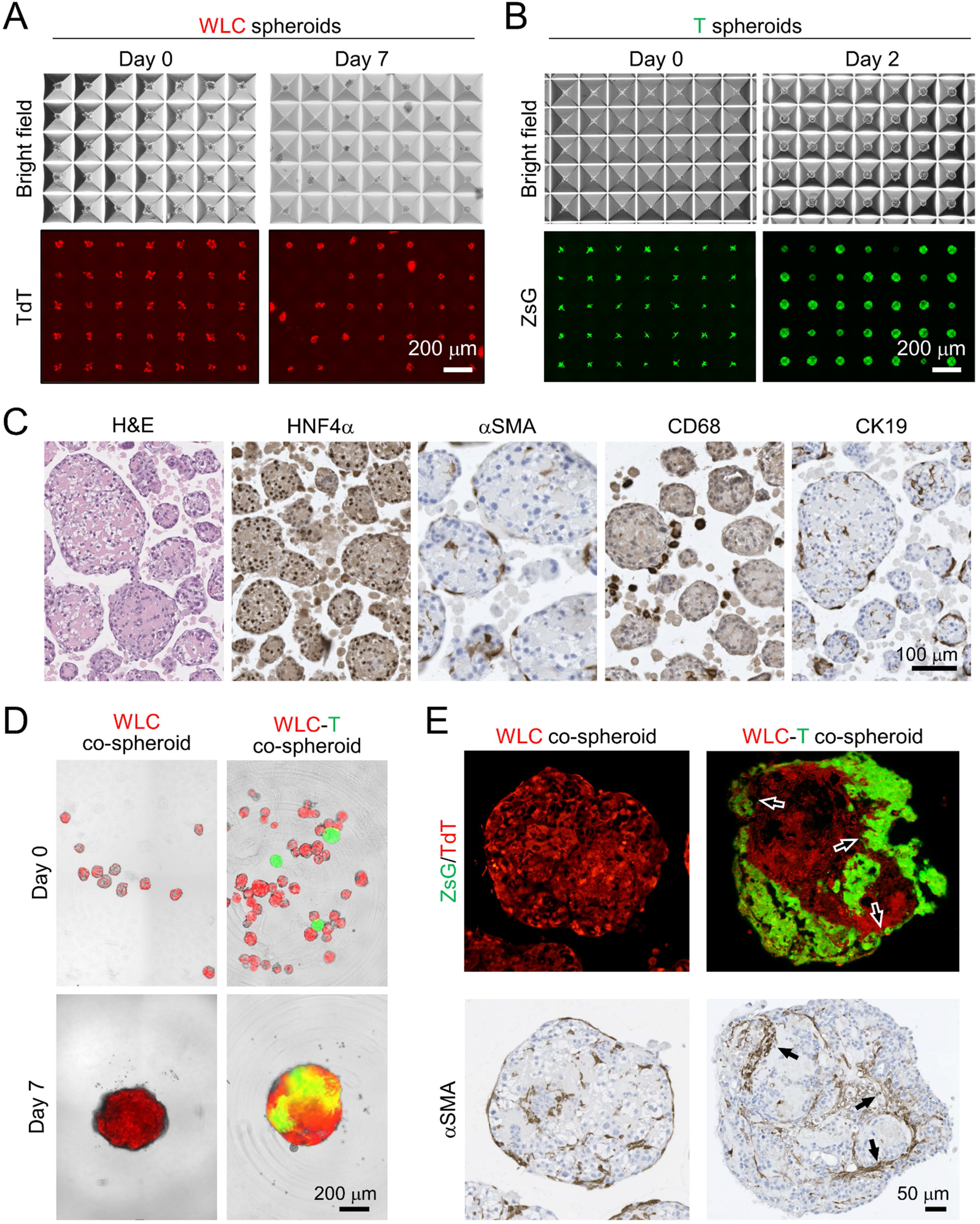
Activation of pMF in the iCCA-liver spheroid coculture. **(A, B)** Generation of the WLC (**A**) and iCCA spheroids (**B**) in AW400 plates. All images share the 200 μm scale bar. **(C)** Histological characterization of the WLC spheroids. Antibodies used are indicated. All images share the same 100 μm scale bar. **(D)** Merged bright-field/ZsG (tumor cells)/TdT (WLC) images of the indicated spheroid cocultures. All images share the same 200 μm scale bar. **(E)** ZsG/TdT fluorescence microscopy and αSMA IHC on the serial sections of WLC and WLC-T co-spheroids collected 7 days after coculturing. All images share the same 50 μm scale bar.

### Peritumoral MFs suppress, and intratumoral MFs promote, iCCA growth in vitro

Since previous studies have consistently shown that aHSCs and CAFs within iCCA tumor mass promote liver tumor growth (6, 7, 20), we examined tumor cell proliferation in our allograft tumors in association with the pMF and iMF content. Surprisingly, we found tumor cells in the pMF^high^ regions at the tumor border in the 1-month allograft tumors had significantly fewer Ki67^+^ cells than those in the tumor core (**Figure 3A** and **3C**). In the 3-month tumors where a large number of iMFs were present in the tumor core, we noticed a strong, positive association between the iMFs content and Ki67 positivity. However, at the invasive tumor border where pMFs accumulated, tumor cell proliferation rate was significantly lower than that of the iMF^high^ tumor core (**Figure 3B** and **3C**). These results suggest that iMFs promote tumor growth as previously reported but pMFs are potentially growth-suppressive. To test this hypothesis, we developed a “2.5-dimensional” (2.5D) coculture system to assess the effect of iMFs and pMFs on iCCA growth. Since aHSCs are the main source of liver MFs, we acquired primary mouse HSCs and induced their activation to MFs via 2D culture on plastic plates (21). Their active proliferation and high expression of αSMA was confirmed via immunofluorescence after three passages in culture (**Suppl Figure S2**). PPTR tumor cells were labeled with CellTracker™ Red CMTPX dye (CellTracker-RFP) which would be diluted into daughter cells upon cell division. Therefore, lower RFP intensity would indicate faster cell proliferation. CellTracker-RFP-labeled PPTR cells were then used to generate iCCA tumor (T)-spheroids, or mixed at 1:1 with MFs to generate T+MF-spheroids. The bottom layer of the 2.5D coculture was prepared by culturing freshly isolated HCs, a 1:1 mixture of HC+MF, or MF-only in 2D condition in 96-well microplates. The bottom layer was allowed to attach for 24 hr and T- or T+MF-spheroids were added on the top at 2-5 spheroids/well. We instantly noticed a slowed expansion of the ZsG^+^ tumor area in the conditions with MFs in the bottom layer (**Figure 4A, a-l**). When T-spheroids were seeded on top of HC+MF mixed at different ratios, we noticed a significant and reversed association between the expansion rate of the ZsG^+^ area and the MF content in the bottom layer (**Suppl Figure S3**). This trend was consistent although showed no statistical significance in the T+MF-spheroid 2.5D cocultures (**Figure 4A, m-x**). Cells in the 2.5D cocultures were collected after four days and examined for their CellTracker-RFP levels. In both T- and T+MF-spheroid cultures, tumor cells placed on top of MFs had the most RFP^+^ cells (**Figure 4B)** and the highest RFP intensity (**Figure 4C)**, indicating their slowest proliferation compared to the other conditions. Tumor cells on top of HC+MF had the second slowest proliferation rate while those on the top of HCs or no bottom layer grew at a similarly high rate (**Figure 4B, C**). Interestingly, when comparing T- and T+MF-spheroids placed on the same bottom layer, tumor cells in the T+MF-spheroids showed consistently and significantly higher proliferation rate indicated by their lower levels of CellTracker-RFP (**Figure 4B, C**). These results are consistent with our observations in the iCCA allograft tumors that there is an intriguing, location-specific effect of MFs on iCCA growth, i.e. iMFs promote tumor growth and pMFs suppress tumor cell proliferation.

**Figure 3.**
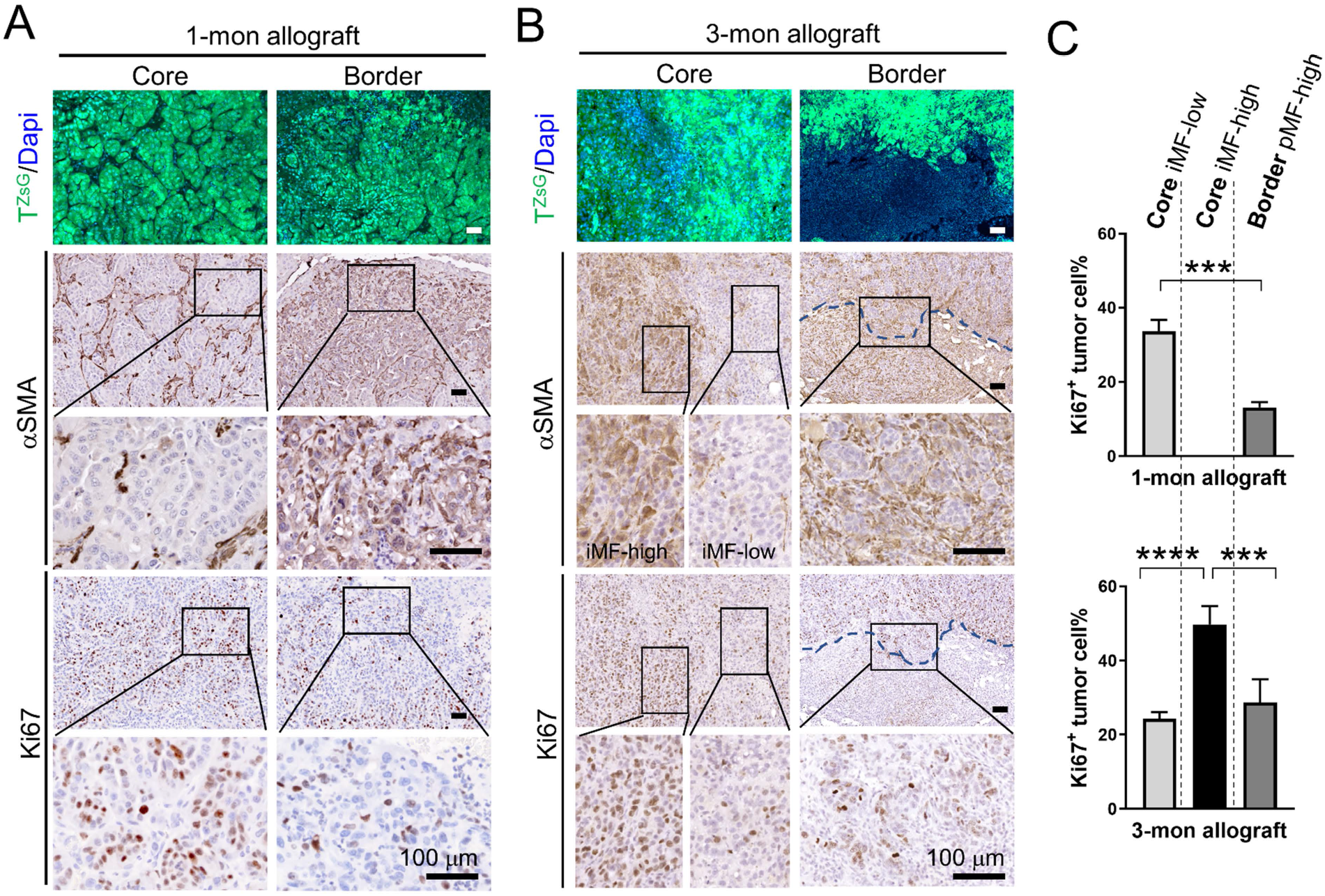
Peritumoral MFs are associated with reduced iCCA proliferation in vivo. **(A, B)** ZsG (tumor cells, T^ZsG^)/Dapi fluorescence microscopy, αSMA and Ki67 IHC in the 1month **(A)** and 3-month **(B)** iCCA allograft tumors. Images in the same row share the scale bar of 100 μm. **(C)** Quantification of Ki67^+^ cells in the indicated areas of the 1- and 3-month iCCA allograft tumors. No MF-high regions present in the core of 1-month tumors. Three tumors were examined at each time point. Student *t*-test, *P* value *** <0.001, ****<0.0001.

**Figure 4.**
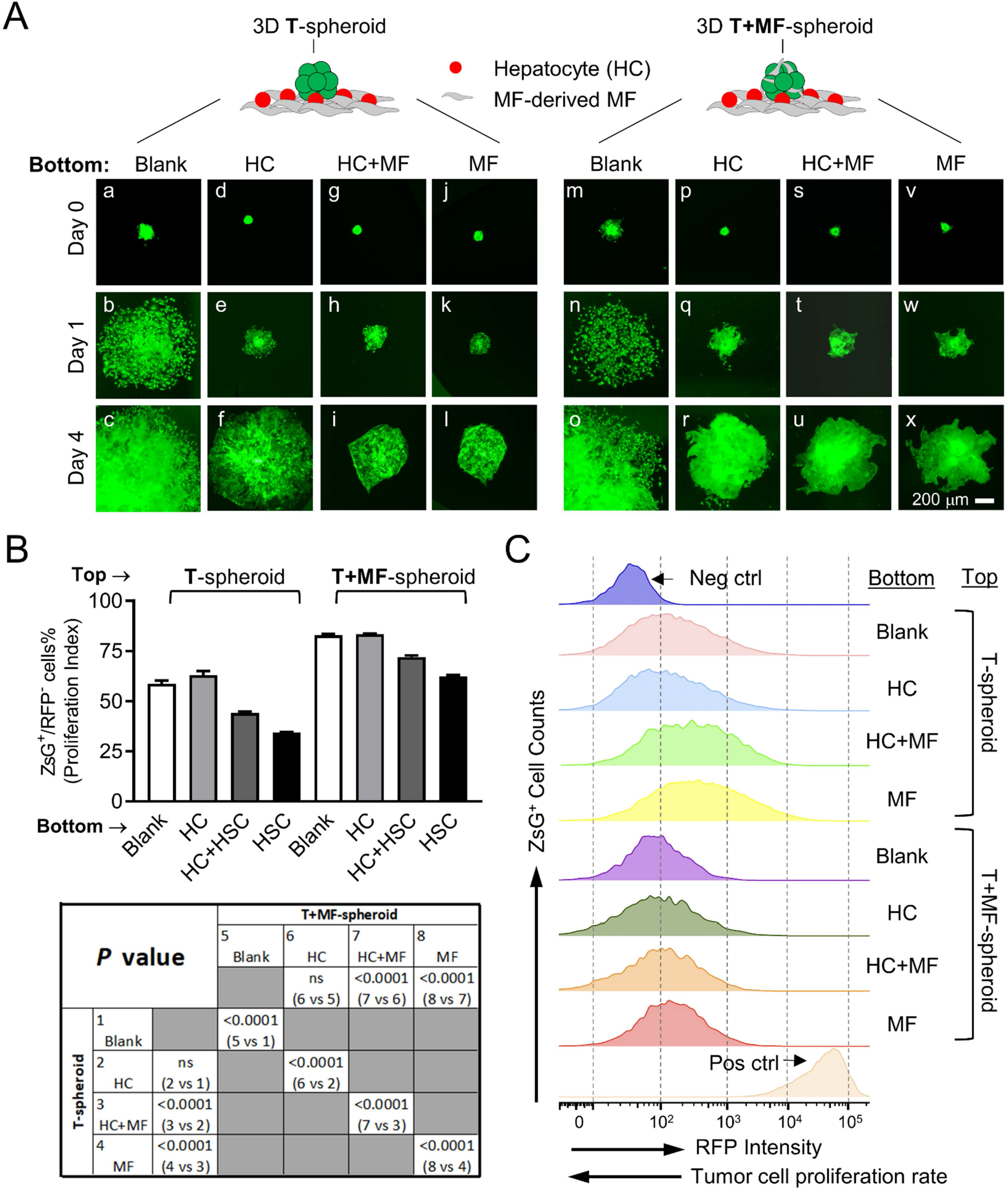
Peritumoral MFs suppress, and intratumoral MFs promote, iCCA growth in vitro. **(A)** Time course images of the ZsG^+^ T- and T+MF-spheroids in the 2.5D coculture with the indicated bottom layers. All images share the same 200 μm scale bar. **(B)** Flow cytometry-based detection of the CellTracker-RFP dye of T- and T+MF-spheroids placed on the indicated bottom layer. Note that RFP-negative cells are plotted to indicate tumor cells proliferation index. *P* values of the Student *t*-test between the indicated groups are shown in the table below. **(C)** RFP intensity histogram of the tumor cells in the indicated groups.

### Elongated iCCA-pMF interaction elicits iCCA cell dissemination in vitro

Since peritumoral HSCs and MFs have been shown to be associated with poor prognosis in HCC patients, we wondered whether the growth-suppressive pMFs might eventually elicit more aggressive behaviors from iCCA tumor cells. To test this, we continued monitoring the T-spheroids 2.5D cocultures up to six weeks (**Figure 5A** and **Suppl Figure S4**). T-spheroids placed on top of MF-containing bottom layers continued growing more slowly than those on the HC-only bottom layer, however, with evident invasion and dissemination from the spheroid surface by two weeks (**Figure 5A, a-h**). Higher MF content on the bottom was associated with longer invasive processes (**Figure 5B**). DTCs were most evident in tumor spheroids placed on top of MF-only (**Figure 5A, h**). Mixing MFs in the tumor spheroids further increased their invasive morphology on the MF-containing bottom layers although no clear increase in tumor cell dissemination was observed (**Figure 5A, i-p**). These observations suggest that pMFs and iMFs play different role in promoting the aggressive behaviors of iCCA tumor cells, that the former works to retard tumor growth but elicit tumor dissemination while iMFs promote tumor growth but play a lesser role in tumor dissemination.

**Figure 5.**
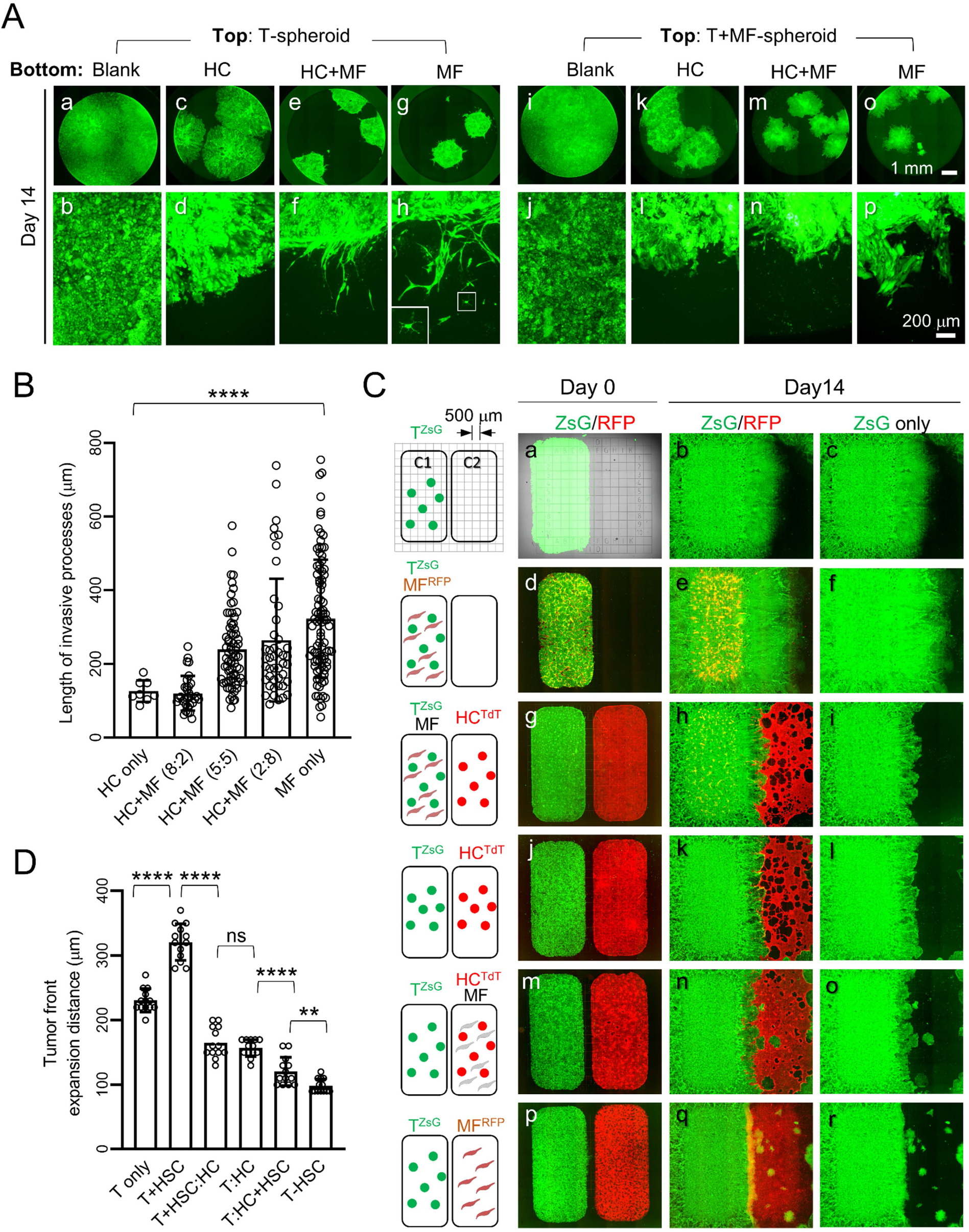
Elongated iCCA-pMF interaction induces tumor cell dissemination. **(A)** ZsG fluorescence images of Day 14 T- and T+MF-spheroid 2.5D cocultures on the indicated bottom layers. Images on the same row share the same scale bar. Inset in **h**: higher magnification image of the boxed disseminated tumor cell. **(B)** Measurement and comparison of the length of the tumor invasive processes on Day 14. Twoway ANOVA comparison, *P* value < 0.0001. **(C)** Day 0 and 14 fluorescence images of the indicated 2D two-chamber cocultures. MFs are labeled with CellTracker-RFP dye in the conditions when no TdT^+^ HCs were added (**d-f** and **pr**). Note that the clonal tumor dissemination only occurs when MFs are present in the C2 chamber (**m-f**). **(D)** Measurement of the advancing distance of the tumor border in the indicated conditions. Student *t*-test, ns, non-significant, *P* value: ns, not significant, ** <0.01, ****<0.0001.

To better visualize the different effect of pMFs and iMFs on tumor cell invasion and dissemination, we developed a 2D coculture system using a two-chamber culture insert to spatially separate tumor cells and MFs (the left and right chamber will be referred to as C1 and C2, respectively) (**Figure 5C**). Culture slides engraved with a 500 μm-grid were used to measure tumor cell movement. Primary HCs were isolated from a *mTmG* mouse and adapted to 2D culture with serial passaging. Different combinations of HCs, MFs, and PPTR tumor cells were seeded as indicated in **Figure 5C**. The insert was removed after 24 hours to allow cells from the two chambers to interact. Similar to our observations from the 2.5D cocultures, iMFs promoted iCCA cell growth in this two-chamber cocultures when tumor cells and MFs were mixed in C1 and C2 was left empty (**Figure 5C, a-c vs. d-f**, and **Figure 5D**). However, this growthpromoting effect of iMFs became less obvious when HCs were placed in C2 (**Figure 5C, g-l** and **Figure 5D**). HCs in C2 significantly slowed the expansion of the tumor compartment although tumor cells were able to steadily push into the HC compartment. Interestingly, we noticed a significantly reduced rate of tumor expansion when MFs were mixed with HCs in C2 as pMFs. However, there was a marked tumor cell invasion and dissemination accompanied with this slowed expansion (**Figure 5C, j-l vs. m-o** and **Figure 5D**). Expansion of the tumor compartment was further decreased when only MFs were placed in C2, and, with further increased tumor cell dissemination (**Figure 5C, p-r**, and **Figure 5D**). Overall, our data support the notion that pMFs suppress the growth of iCCA tumor cell but promote tumor cell invasion and dissemination in vitro.

### Peritumoral MFs induce Vcam1 upregulation in iCCA tumor cells

Next, we investigated the potential molecular mechanisms involved in iCCA-pMF interaction. Since cytokines are known key players in fibroblast-associated tumor invasion and metastasis (22, 23), we performed a cytokine array assay using conditioned media (CM) collected from 2.5D cocultures of tumor spheroids on top of HC (T:HC) and on top of MFs (T:MF) and their mono cultures as controls. We used CMs from Day 4 when the growth suppressive effect of pMFs was evident. We noticed a mild but significant increase in Vascular Cell Adhesion Molecule 1 (Vcam1) in T:MF CM compared to the others (**Figure 6A**). and we have previously reported that Vcam1 levels were increased in metastatic tumors in PPTR genetic and allograft models (16). Via flow cytometry, we confirmed an increase in Vcam1^+^ tumor cells in T:MF coculture on Day 4 (**Figure 6B**). Surprisingly, although Vcam1 has been shown to promote solid tumor metastasis (24, 25), we noticed that Vcam1 levels dropped significantly in the tumor cells collected from the six-week T:MF coculture when many tumor cells had loosely detached from the spheroid (**Figure 6C**, and **Suppl Figure S4**). In the iCCA allograft tumors, we found Vcam1 upregulation in the tumor cells in contact with pMFs in early-stage 1-month tumors while little Vcam1 expression was seen in the tumor core (**Figure 6D, a-e**). Vcam1 expression became more heterogeneous in 3-month allograft tumors. High Vcam1 expression was mostly found in the tumor cells at the invasive borders where pMFs accumulated, while no such association was seen between Vcam1 expression and iMFs within the tumor core (**Figure 6D, f-i**). Interestingly, we found ZsG^+^ DTCs in both liver and lung from our allograft model were consistently Vcam1-negative while metastatic tumors had high levels of Vcam1 (**Figure 6E**). These data suggest that Vcam1 is likely important for tumor establishment in a suppressive environment consisting of pMFs and others. But its function as an adhesion molecular may have a negative impact on tumor cell dissemination.

**Figure 6.**
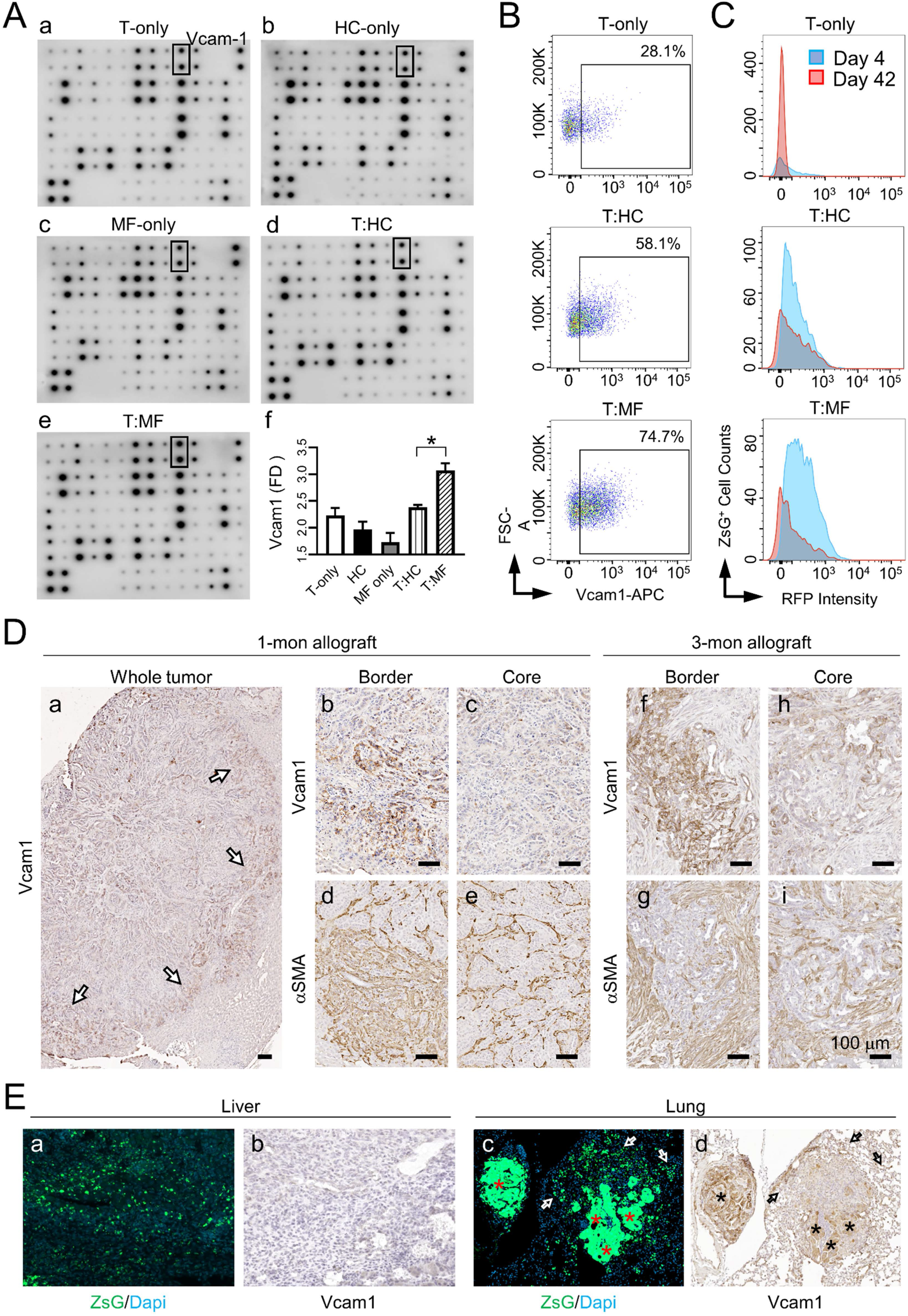
iCCA-pMF interaction leads to dynamic changes in Vcam1 expression in tumor cells. **(A)** Cytokine array assays using the CM from the indicated cultures on Day 4 detected an upregulation of Vcam1 in T (top)+MF (bottom) (T:MF) 2.5D coculture. **f:** quantification of the Vcam1 levels in the CM from the indicated cultures. Student *t*-test, *P* value * <0.05. **(B)** Vcam1 detection in the indicated 2.5D cocultures on Day 4 via flow cytometry. The percentage of Vcam1^+^ tumor cells in each condition as indicated and T:MF coculture has the most Vcam1^+^ tumor cells. **(C)** Histogram of Vcam1 level in the tumor cells from the indicated cocultures on Day 4 and 42, showing a decrease in Vcam1 levels in Day 42 cultures. **(D)** Vcam1 and αSMA IHC on the serial sections of the indicated allograft tumors. All scale bars are 100 μm. Arrows in **(a)**: Vcam1^high^ cells at the tumor border. **(E)** ZsG/Dapi fluorescence and Vcam1 IHC on the serial sections of the live (**a**, **b**) and lung (**c, d**) from iCCA allografts showing decreased Vcam1 levels in the DTCs. Asterisks in (**c, d**): Vcam1^high^ tumor cells within the lung metastases; arrows in (**c**, **d**): Vcam1^low^ DTCs.

### Targeting Vcam1 in iCCA transplantation models promote tumor metastasis

Next, we examined the importance of Vcam1 in tumor growth and metastasis in our iCCA models. We were not able to directly manipulate Vcam1 expression in PPTR tumor cells because these cells had shown strong resistance to transduction. A small subpopulation of PPTR cells that we were able to transduce with a control vector produced tumors with different histopathology (data not shown). Therefore, we chose to use a blocking strategy to examine the effect of Vcam1 on iCCA growth and dissemination in vitro and in vivo. We found PPTR tumor cells treated with a reported Vcam1 neutralizing antibody (Vcam1^Ab^) (26) showed increased, not decreased, migration ability (**Supple Figure S5A**). We did not observe any changes in PPTR tumor cell proliferation after Vcam1^Ab^ treatment in vitro (**Supple Figure S5B**). We also did not find a positive association between Vcam1 expression and tumor cell proliferation in the allograft tumors (**Supple Figure S5C**). In the orthotopic model treated with Vcam 1^Ab^ for three weeks, we found tumors at the injection site were smaller than those in the IgG-treated mice (**Figure 7A, a-d**). However, Vcam1^Ab^**-**treated mice had developed visible micrometastases in the liver while the IgG-treated mice had not (n=3 per group) (**Figure 7A, b**). Small disseminating tumor clones were evident on the border of Vcam1^Ab^-treated tumors but not the IgG-treated tumors (**Figure 7A, e, f**). No overt lung metastases had developed in either groups at this stage. However, there were consistently more ZsG^+^ DTCs in the lungs of Vcam1^Ab^**-**treated mice than the IgG-treated ones (**Figure 7A, g-h**). After six weeks of treatment, the different in tumor size was less evident between the two groups. However, Vcam1^Ab^ group consistently had more metastases in both liver and lung than the IgG group (n=3 per group) (**Figure 7B**). Metastases were present in the non-injected liver lobes in the Vcam1^Ab^ group while the IgG group had DTCs in the liver but no overt metastases (**Figure 7B, a-d**). Lung metastases were found in both groups but consistently more in the Vcam1^Ab^ group (**Figure 7B, eh**).

**Figure 7.**
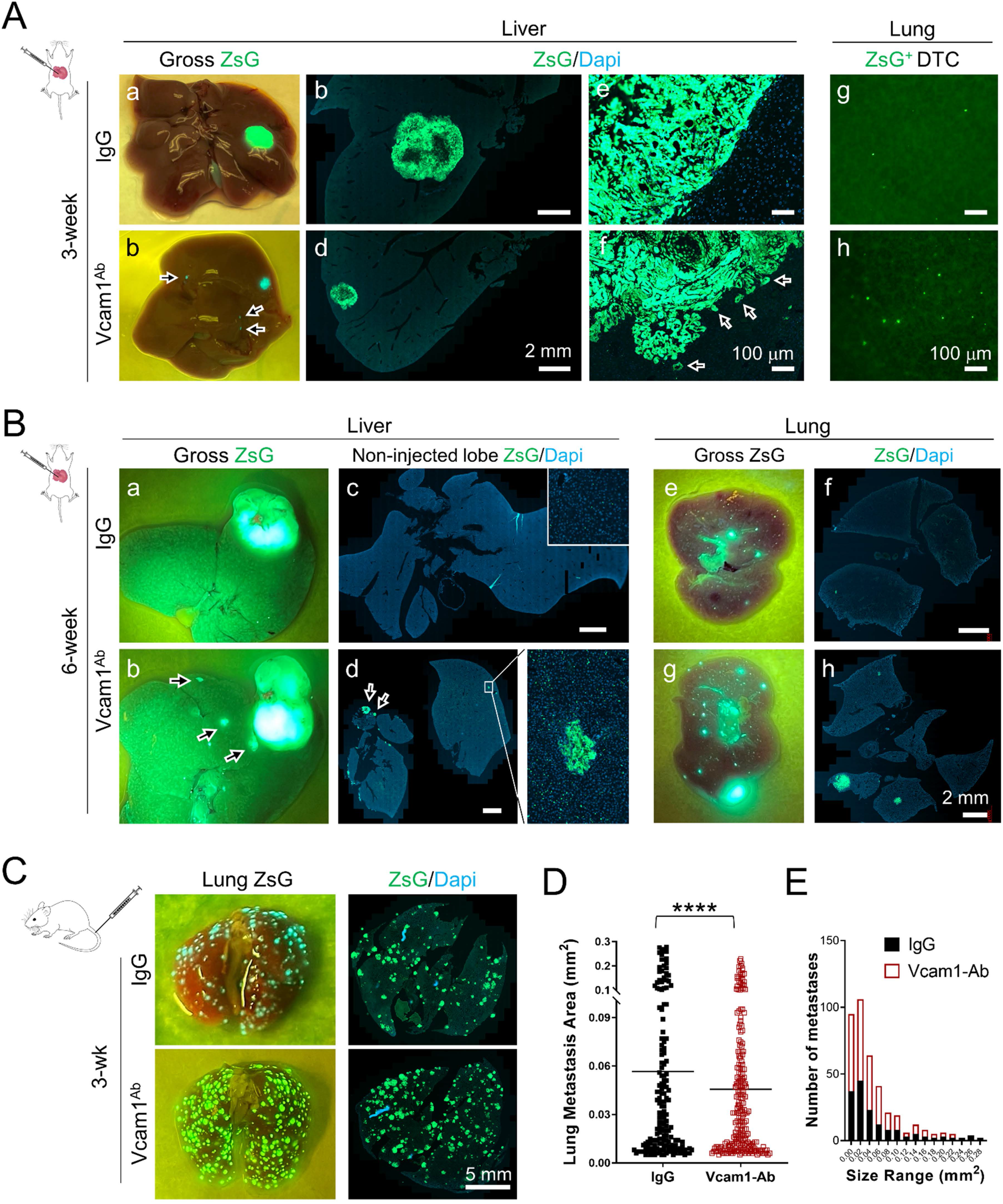
Vcam1 blocking promotes tumor metastasis in the iCCA transplantation models. **(A)** Gross ZsG and ZsG/Dapi images of the liver and lung collected from the orthotopic models after three weeks of IgG or Vcam1^Ab^ treatment. **(a, b)** Gross live ZsG showing the smaller tumor in the Vcam1^Ab^ -treated liver; arrows in **(b)**: micrometastases in the Vcam1^Ab^ -treated liver. **(b-f)** ZsG/Dapi fluorescence images of liver sections; arrows in **(f)**: small disseminating clones on the border of Vcam1^Ab^ -treated tumor. **(g, h)** ZsG fluorescence of the lung lobes showing a higher number of DTCs entering the Vcam1^Ab^ -treated lung. **(B)** Similar gross ZsG and ZsG/Dapi images of the liver and lung collected from the orthotopic models after six weeks of treatment showing increased liver and lung metastasis in the Vcam1^Ab^ - treated mice. Arrows in **(b d)**: metastases in the Vcam1^Ab^ -treated liver. **(C)** Gross ZsG and ZsG/Dapi images of the lung metastases collected from the TVI model treated with IgG or Vcam1^Ab^ for three weeks showing increased metastasis in Vcam1^Ab^ -treated mice. **(D)** Quantification of the area of individual lung metastases areas from the TVI model showing larger lung metastases in the IgG-treated mice. Student *t*-test, *P* value ****<0.0001. **(E)** Histogram of the size distribution of the lung metastases from the TVI model showing the larger number of small lung metastases in the Vcam1^Ab^ -treated mice.

To further validate the metastasis-promoting effect of Vcam1 neutralization in vivo, we utilized an iCCA lung metastasis model established via tail vein injection (TVI) of PPTR tumor cells. Although no liver pMF-tumor interaction was involved in this model, we have shown that lung metastases in the orthotopic model expressed high levels of Vcam1 (Figure 6D). Therefore, this TVI metastasis model would allow us to evaluate the importance of Vcam1 to iCCA lung metastasis and its potential therapeutic value in metastatic iCCA. Similar to the orthotopic model, we found the TVI group treated with Vcam1^Ab^ for three weeks, instead of having suppressed lung metastasis, developed more lung metastases than those treated with IgG (n=5 per group) (Figure 7C and Suppl Figure S6). Interestingly, we found the size of the lung metastases in the IgG-treated mice was larger on average than those in the Vcam1^Ab^-treated mice, while the latter developed more metastases overall (Figure 7, D&E). These findings from both the orthotopic and TVI transplantation models suggest that Vcam1 plays an important role in supporting iCCA tumor growth in the liver and lung but reduced Vcam1 activity facilitates tumor cells to successfully disseminate from the established tumor mass. Therefore, similar to MFs, Vcam1 in iCCA is also beyond simple pro- or anti-tumorigenic and plays dynamic roles in tumor progression and dissemination.

## DISCUSSION

Using an orthotopic model of metastatic iCCA and multiple novel multi-lineage cellular models we established in the laboratory, we showed that there was an interesting MF dynamics during iCCA development. At the early stage of iCCA tumorigenesis, pMFs rapidly accumulate at the tumor border. When tumor gets larger, there is an increase in iMFs within the tumor while pMFs become particularly enriched at the tumor invasive front. We found iMFs and pMFs are different in their roles in promoting iCCA aggressiveness. Intratumoral iMFs are growth-promoting as previously reported (6, 7), while pMFs exhibit a surprising suppressive effect on iCCA growth. However, we found prolonged iCCA-pMF interaction elicits tumor cell dissemination, which is in line with the previous observations in HCC patients that activated HSCs and MFs in the peritumoral liver are associated with tumor metastasis and relapse (11, 12).

Based on our study, we propose that the initial MF activation and accumulation at the tumor border is a defensive response of the liver to iCCA tumorigenesis, intended to restrain local invasive tumor growth and, in turn, preserve liver function. Indeed, recent studies have started to argue that cirrhosis, a late-stage liver fibrosis condition that heavily involves MF activation, may be a liver-protective response rather than a risk factor for live cancer (27). However, our study finds that the growth suppression by pMF has an adverse consequence which is the eventual elicitation of more aggressive behaviors from tumor cells leading to tumor dissemination. This observation is in line with the role of cell competition in selecting aggressive cancer behaviors (28). iCCA and many aggressive solid tumors are known to be highly heterogeneous and plastic, and tumor cells can readily change their behaviors depending on the environmental cues they receive. Tumor suppression by pMFs may lead to a selection process for aggressive iCCA cells while attempting to contain tumor growth (29). Although we lack the tools to efficiently detect tumor dissemination in iCCA patients as we do for our mouse models, the high rates of tumor relapse in iCCA patients, even after curative resection, suggests that tumor dissemination is common in these patients. This could potentially be elicited by the large pMF population around iCCA tumors as we have shown in this study. Based on our findings, we suspect that MF depletion at the early stage of tumorigenesis would potentially dampen liver’s ability to suppress tumor growth and would lead to accelerated tumor development. However, with the continuous increase in protumorigenic iMFs and weakened ability of pMFs to restrain aggressive tumor growth, MF depletion in late-stage tumors may in fact an effective strategy to slow tumor progression. We suspect that the different timing of MF depletion may account for, at least partially, to the contradicting tumor outcomes in the previously reported CAF-depletion studies (7, 30–33). A genetic approach capable of temporal- and spatial-specific depletion of MFs, when it becomes available, would be the most definitive in defining their dynamic role in iCCA development. Single-cell transcriptomic analyses using our coculture systems and mouse models will also be necessary to pinpoint the molecular mechanisms behind the differential roles of iMFs and pMFs play in iCCA development.

Since the role of intratumoral fibroblasts have been widely studied for liver cancer, we focused on pMF-tumor interaction in this study for molecular investigations. We show that Vcam1 is upregulated in the tumor cells at the early phase of pMF-iCCA interaction. But Vcam1 level drops when tumor cells start to disseminate. Although Vcam1 has been previously shown to promote metastasis (24, 25), our tracking of Vcam1 expression in the early- and late-stage metastatic iCCA allograft tumors suggests that its function is beyond pro- or anti-metastatic. Vcam1 facilitates tumor growth in both liver and lung. But it negatively impacts tumor cell dissemination as Vcam1 blocking in our mouse models increases metastasis in both liver and lung. We need to acknowledge that the PPTR tumor cells we used in this study are highly metastatic. It is possible that their dependency on Vcam1 is different from tumor cells with low metastatic potential. We also acknowledge that our strategy of using a Vcam1 neutralizing antibody, due to the resistance of PPTR cells to genetic manipulation, only allows us to study the systemic effect of Vcam1 in iCCA metastasis. Such an approach does not differentiate the contributions from other Vcam1-expressing cells to the metastasis in our models. Although Vcam1 is predominantly expressed in the tumor cells in our mouse models, it does not rule out the possibility that low Vcam1 activity from other nonmalignant cell types may be critical to iCCA metastasis as reported in other solid tumors (24, 25). However, our study does argue against the previous proposals of targeting VCAM1 as a therapeutic strategy to treat metastatic disease (34). Our results clearly indicate that highly aggressive tumors could benefit from Vcam1 blocking to disseminate more efficiently. A more detailed understanding of the dynamic involvement of Vcam1 in the complex metastasis cascade is needed before we can adequately assess its therapeutic value in metastatic cancer.

Overall, our study suggests that iCCA-MF interaction, or tumor-host interaction in a larger picture, is beyond simple pro- or anti-tumorigenic. We theorize that iCCA metastasis is an ongoing competitive process between the attempt of pMFs to block local growth of the tumor and that of tumor cells to break through the suppression. Future studies are in line to identify specific mechanisms underlying this complex tumor-host crosstalk with a goal to simultaneously address local tumor growth and early steps in metastatic progression.

## METHODS

### Mice

Two-month-old male and female *B6.129(Cg)-Gt(ROSA)26Sor^tm4(ACTB-tdTomato,-EGFP)Luo^/J (mTmG)* mice (Stock No. 007576, Jackson Laboratory, Bar Harbor, ME, USA) were used for primary cell isolation from the liver. Two-month-old male and female *Crl:CD1-Foxn1^nu/nu^* (CD-1 nude) mice (Strain Code 086, Charles River, Wilmington, MA, USA) were used for iCCA orthotopic transplantation and tail vein injection. Animal protocols were approved by the St. Jude Animal Care and Use Committee. All mice were maintained in the Animal Resource Center at St. Jude Children’s Research Hospital (St. Jude). Mice were housed in ventilated, temperature- and humidity-controlled cages under a12-hr light/12-hr dark cycle and given a standard diet and water *ad libitum.*

### Cell culture

Two iCCA tumor cell lines, PPTR-1 and −2, were established from **P*rom1^CreERT2^; <u>p</u>ten^flx/flx^; <u>T</u>p53^flx/flx^ <u>R</u>osa-ZsGreen* (PPTR) liver cancer organoids and maintained in DMEM (Corning Inc, Corning, NY, USA) supplemented with 10% fetal bovine serum (FBS) and antibotics (16). Primary mouse HSCs were purchased (Cat No. M5300-57, ScienCell Research Laboratories, San Diego, CA, USA) and cultured according to the manufacturer’s instruction. All iCCA-liver coculture was maintained in a 1:1 mixture of mouse HepatiCult™ Organoid Growth Medium (STEMCELL) and previously reported cholangiocyte organoid culture medium supplemented with 1% GFR matrigel was used in all cocultures (35). AggreWell 400 (AW400) culture plates (STEMCELL Technologies, Vancouver, Canada) were used to generate iCCA and liver spheroids. Two-chamber culture insert and μ-slide 8-well Grid-500 culture slides from ibidi (Gräfelfing, Germany) were used in 2D coculture. See Supplementary Material for details.

### iCCA orthotopic and tail vein transplantation models

For iCCA orthotopic transplantation, PPTR-1 tumor cells were surgically injected into the liver of two-month-old male and female CD-1 nude mice at 1× 10^5^/mouse in 4 μl cold growth factor-reduced (GFR) matrigel (Corning). For tail vein injection (TVI)-based iCCA lung metastasis model, PPTR-1 tumor cells were injected into two-month-old CD-1 nude mice at 1×10^6^/mouse in 100 μl PBS via TVI.

### IgG and Vcam1 antibody treatment

For in vivo assays, mice were treated three days after orthotopic or TVI transplantation with *InVivoMAb* anti-mouse CD106 (Vcam1) antibody (Vcam1^Ab^) (Bioxcell, Lebanon, NH, USA) or rat IgG (R&D System, Minneapolis, MN, USA) at 0.25 mg/mice via TVI for 3 weeks, twice a week, five mice/group. For in vitro assays, cultured cells were treated with 10 μM Vcam1^Ab^ or IgG for 4 days.

### Reagents for cell labeling and detection

All flow cytometric analyses were performed on a BD LSRFortessa™ cell analyzer (BD Biosciences) and Flow data were analyzed using FlowJo_V10. See Supplementary Material for details.

Vcam1 detection: APC anti-mouse CD106 (Vcam1) antibody (BioLegend, San Diego, CA, USA). Cell proliferation: CellTracker™ Red CMTPX dye (Invitrogen, Carlsbad, CA, USA).

### Cytokine array assay

Conditioned media (CM) were collected from cell cultures after four days. The levels of secreted cytokines in the CM were measured using Mouse Cytokine Antibody Array 3 (RayBiotech, GA, USA) according to the manufacturer’s instructions.

### Immunohistochemistry and immunofluorescence

Paraffin sections (4 μm) of liver spheroids and tissues were prepared by HistoWiz Inc. (Brooklyn, NY) and analyzed by direct fluorescence microscopy, H&E staining, and IHC. Direct ZsG and TdT fluorescence images were taken from the sections prior to IHC. Primary HSCs cultured on glass chamber were subject to standard immunofluorescence. Antibodies used included anti-Ki67 (Abcam, Cambridge, MA, USA, ab16667, 1:200); anti-CK19 (Abcam, ab133496, 1:250); anti-Vimentin (Abcam, ab92547, 1:500); anti-HNF4a (R&D System, Minneapolis, MN, USA, PP-K9218-00, 1:125); anti-Cyp3A (Abcam, ab3572, 1:500); anti-aSMA (Abcam, ab124964, 1:1000); Vcam1 (Abcam, ab134047, 1:1000) and anti-CD68 (Abcam, ab125212, 1:1000).

### Microscopy, Image-based Quantification and Statistical Analysis

All cocultures were monitored daily by using an ECLIPSE Ts2R fluorescence microscope (Nikon, Minato City, Tokyo, Japan) and Lionheart FX Automated Microscope (Winooski, VT, USA). The ZsG^+^ tumor cell area and TdT^+^ HC area in cocultures were measured by Image J and plotted in GraphPad Prism 7. See Supplementary Material for details. Two-tailed student *t* test was performed in GraphPad Prism 7 to compare two independent pairs of groups. One-way ANOVA was performed when two or more groups were compared. *P* value ≤0.05 was considered statistically significant.

### Study approval

Animal experiments were approved by St. Jude Animal Care and Use Committee. The de-identified iCCA patient tumor samples were obtained under a protocol approved by the Institutional Review Boards at both St. Jude Children’s Research Hospital and The University of Tennessee Health Science Center.

## Abbreviations

ICC: Intrahepatic cholangiocarcinoma
ECC: extrahepatic cholangiocarcinoma
HCC: Hepatocellular carcinoma
CAF: Cancer-associated fibroblast
HSC: Hepatic stellate cells
aHSC: activated HSCs
MF: myofibroblast
pMF: peritumoral myofibroblast
iMF: intratumoral myofibroblast
HC: Hepatocyte
WLC: Whole liver cell
DTC: Disseminated tumor cells
PPTR: Prom1^CreERT2^; Pten^flx/flx^; Tp53f Rosa-ZsGreen
mTmG: B6.129(Cg)-Gt(ROSA)26Sor^tm4(ACTB-tdTomato,-EGFP)Luo^/J
ZsG: Rosa-ZsGreen
TdT: Rosa-tdTomato
RFP: Red Fluorescent Protein
GFP: Green Fluorescent Protein
AW400: AggreWell 400
CM: Conditioned Media
TVI: Tail Vein Injection

