## Supplementary figures and methods for "Paradoxical Roles of Peritumoral Myofibroblasts in Intrahepatic Cholangiocarcinoma Growth and Metastasis"

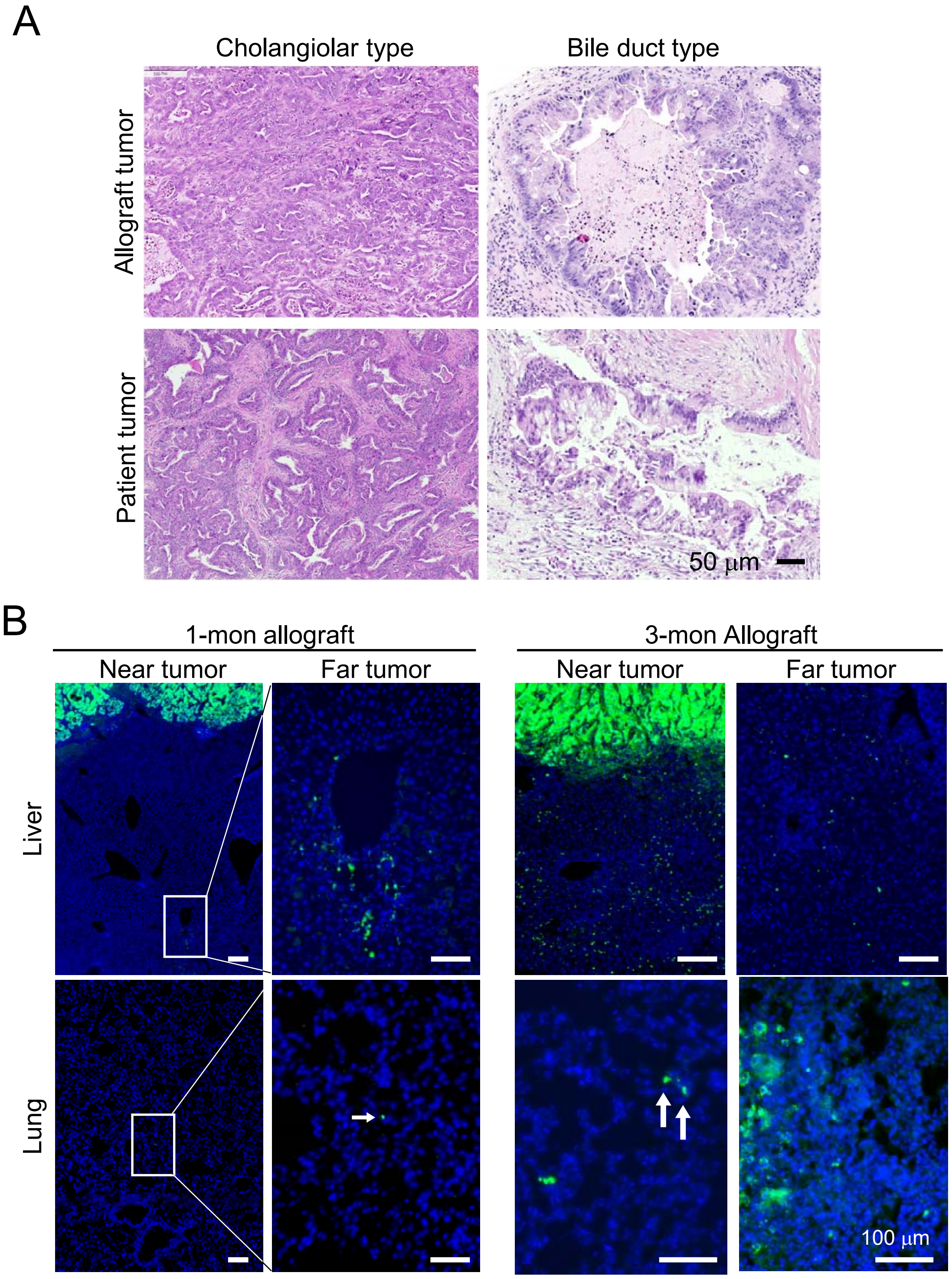


**Supplemental Figure S1. Generation of a metastatic iCCA orthotopic allograft model.**

**(A)** H&E images of the allograft and patient iCCA tumors showing the two types of characteristic histology. All images share the same 50 μm scale bar. **(B)** Dissemination of the $ZsG^{+}$ tumor cells in the liver and lung of the 1- and 3-month iCCA allograft tumors. All scale bars are 100 μm.

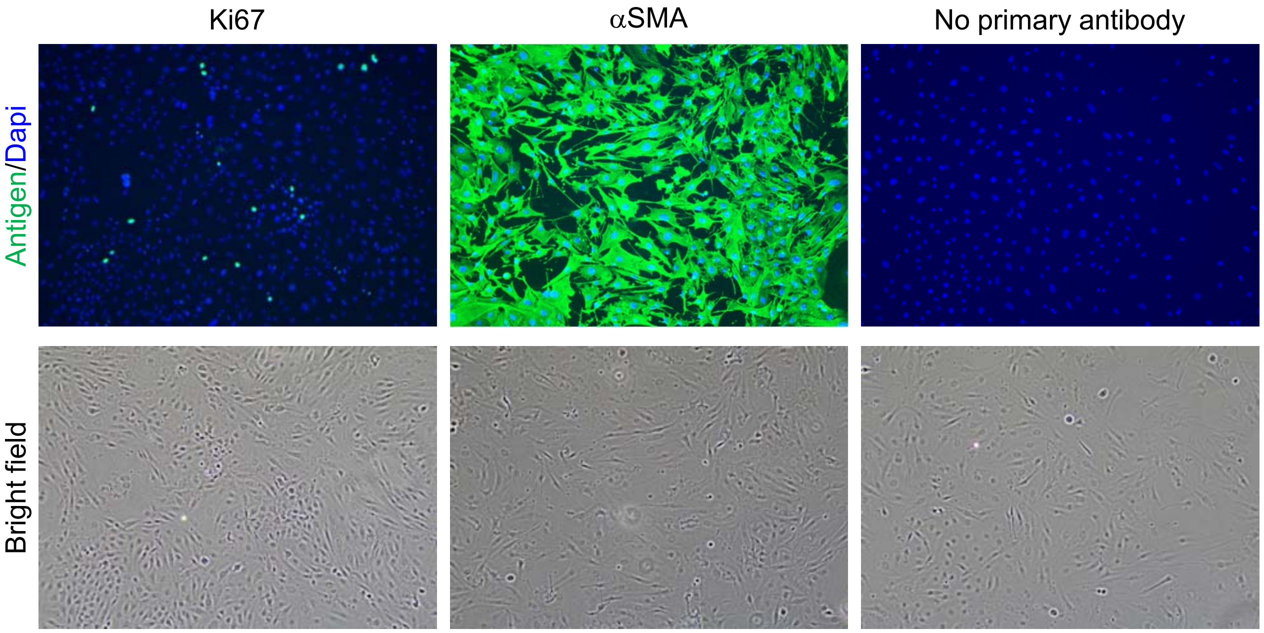


**Supplemental Figure S2. Activation of mouse primary HSCs in 2D culture.**

Ki67 and αSMA immunofluorescence staining of the mouse primary HSCs after three passages on 2D plastic plates, showing their active proliferation and strong expression of the MF marker αSMA.

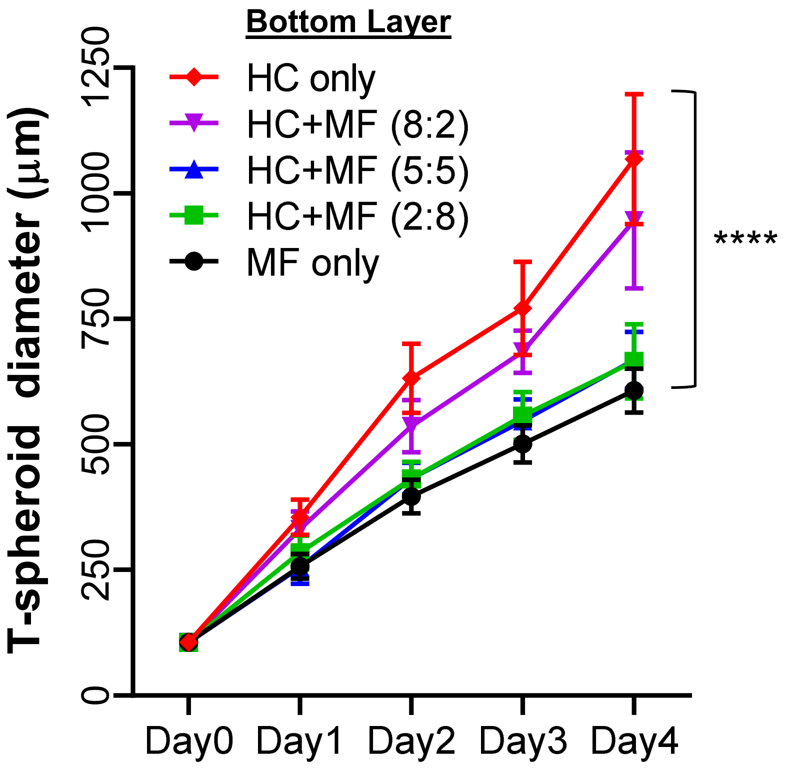


**Supplemental Figure S3. Peritumoral MFs retard iCCA spheroid expansion in the 2.5D cocultures.**

Measurement and comparison of the diameters of the iCCA spheroids show decreased tumor expansion rate with increased MF content in the bottom layer. Two-way ANOVA comparison, $P$ value < 0.0001.

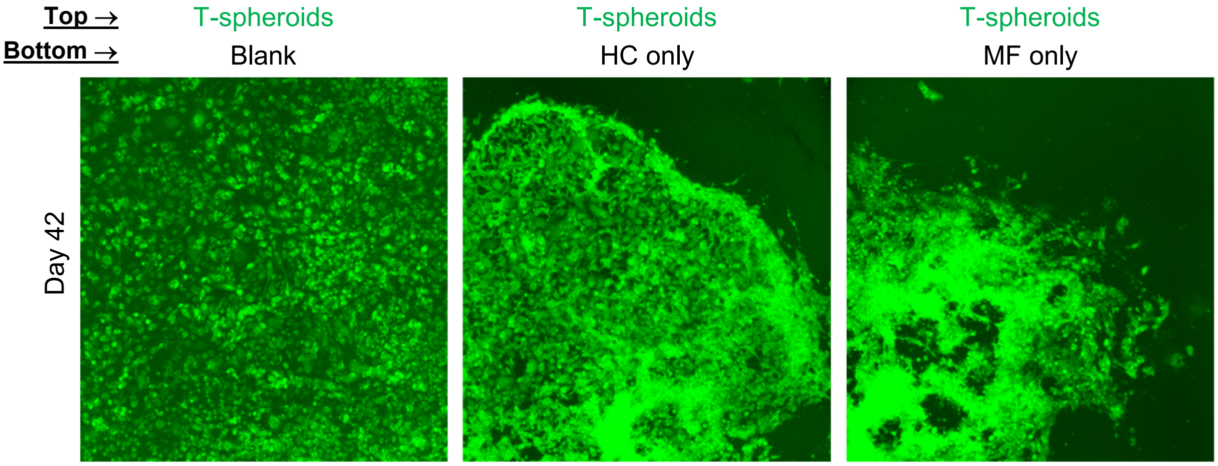


**Supplemental Figure S4. Tumor spheroids in the long-term iCCA 2.5D culture.**

ZsG fluorescence images of the iCCA T-spheroids cultured without a bottom layer (blank), on top of HCs only or MFs only on Day 42.

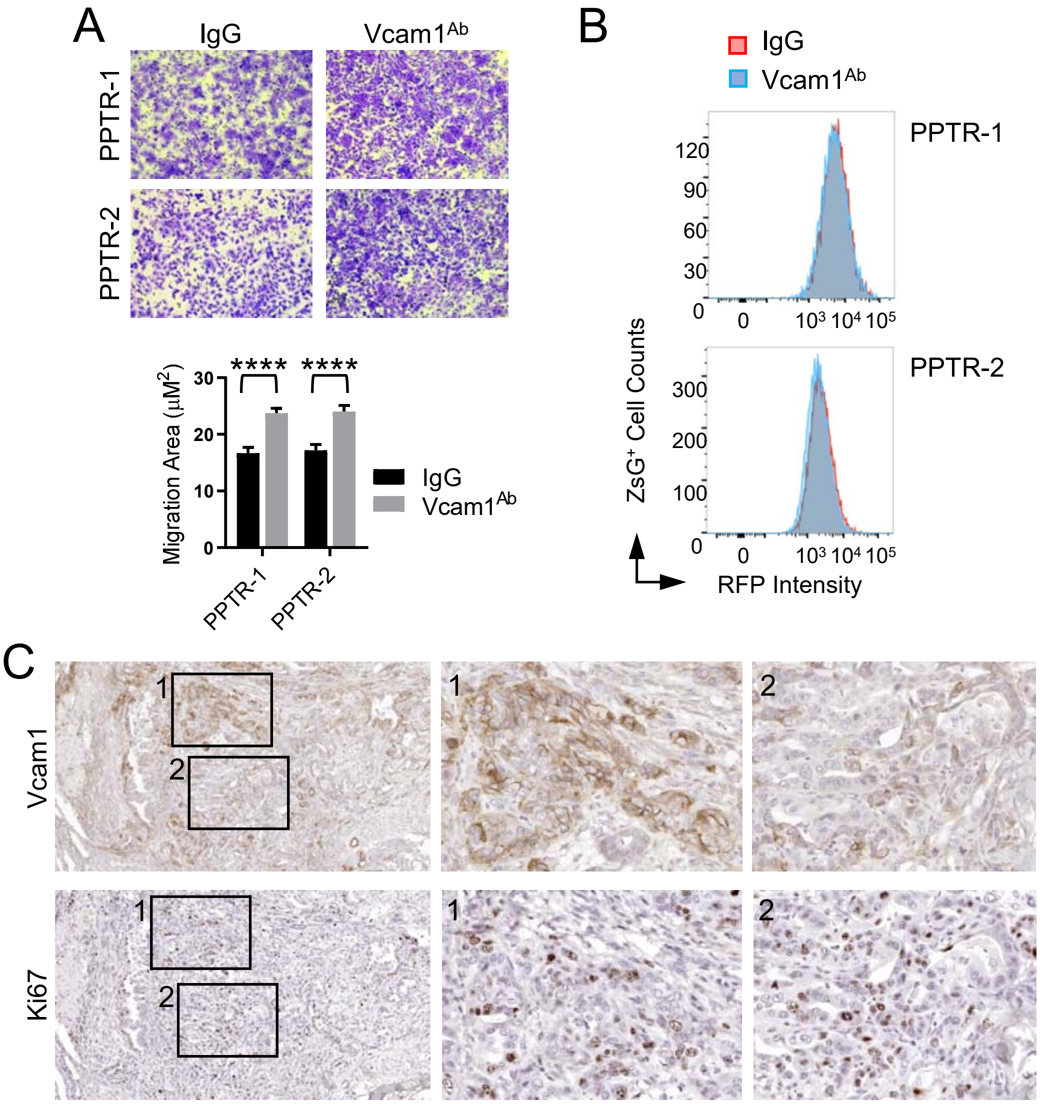


**Supplemental Figure S5. Vcam1 is associated with iCCA tumor cell migration but not proliferation.**

**(A)** Transwell migration assay and quantification of the PPTR cells treated with IgG or $Vcam1^{Ab}$ showing increased cell migration with Vcam1 blocking. Student *t*-test, *P* value <0.0001. Cells derived from two PPTR tumors were tested. **(B)** Flow cytometry of CellTracker proliferation assay of the PPTR tumor cells treated with IgG or $Vcam1^{Ab}$ showing no changes in cell proliferation with Vcam1 blocking. **(C)** Vcam1 and Ki67 IHC on serial sections of a 3-month iCCA allograft tumor. No association was found between Vcam1 expression and Ki67 positivity.

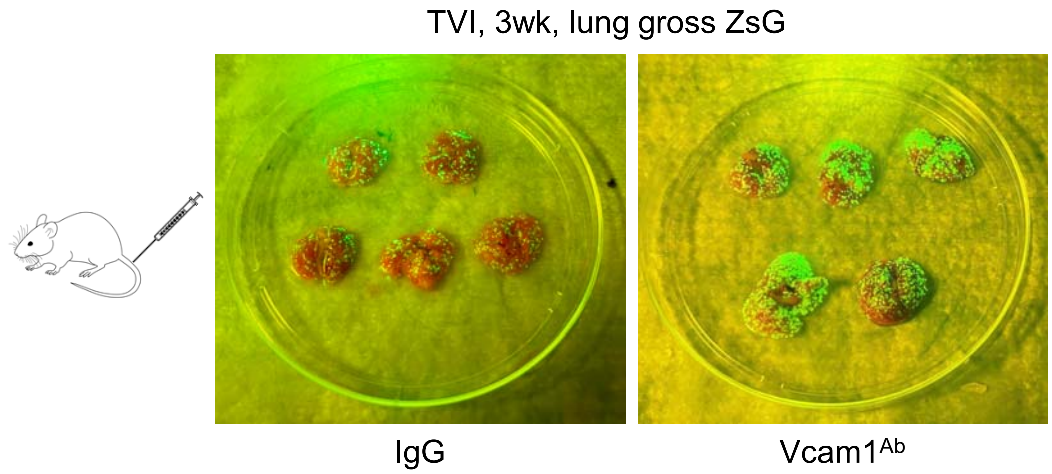


**Supplemental Figure S6. Vcam1 blocking promotes iCCA lung metastasis in the TVI model.**

Gross ZsG images of the lungs collected from the iCCA TVI model treated with IgG or Vcam1$^{Ab}$ for three weeks. Lungs from the Vcam1$^{Ab}$ group consistently have more lung metastases.

**SUPPLEMENTARY MATERIAL**

### Cell Culture

#### 1. Primary liver cell isolation

Liver cells were isolated from two-month-old *mTmG* mouse liver using the standard two-step collagenase perfusion method.[1, 2] Briefly, mice were anesthetized with isoflurane, the liver was perfused with 30 ml of prewarmed EGTA solution (0.5 mM) through the inferior vena cava for 10 min, and then perfused with 30 ml of prewarmed pronase solution (14 mg/mouse; Sigma-Aldrich, St. Louis, MO, USA) followed by 30 ml of collagenase solution (3.7 U/mouse; Roche, Basel, Switzerland) for 10 min each. The liver was then removed and transferred to a sterile Petri dish and gently minced with forceps. The liver was further digested with the pronase/collagenase solution with 1% DNase I (Sigma-Aldrich) for 25 min at 40 °C. Cells were filtered by a 70-μm filter and used directly as unfractionated WLCs, or centrifugated at 50 ×*g* for 3 min to pellet HCs. WLCs and HCs were washed twice in cold Advanced DMEM/F12 (Thermo Fisher Scientific, Waltham, MA, USA) and subjected to spheroid culture.

#### 2. Mouse hepatic stellate cell culture

Primary mouse HSCs were purchased (Cat No. M5300-57, ScienCell Research Laboratories, San Diego, CA, USA) and cultured in Stellate Cell Medium (Cat No. 5301, ScienCell) supplemented with 1% stellate cell growth supplement (Cat No. 5352, ScienCell), 2% FBS, and 1% Penicillin/Streptomycin Solution.

#### 3. Spheroid culture

Freshly isolated WLCs and HCs were seeded in 24-well AggreWell 400 (AW400) culture plates (STEMCELL Technologies, Vancouver, Canada) at 125 cells/microwell, and cultured for three days under continuous shaking at 80 rpm on a $CO_2$-resistant Orbital shaker (Thermo Fisher Scientific, Waltham, MA, USA). A 1:1 mixture of mouse HepatiCult™ Organoid Growth Medium

(STEMCELL) and previously reported cholangiocyte organoid culture medium[3] supplemented with 1% GFR matrigel was used to culture all spheroids. The cholangiocyte organoid culture medium was prepared by supplementing advanced DMEM/F12 (Thermo Fisher) with 50% conditioned medium from L-WRN cells (ATCC CRL-3276™, Manassas, VA, US), 10 mM HEPES, 1% GlutaMax, 1% Penicillin-Streptomycin, 2% B27 (Thermo Fisher), 1% N2 (Thermo Fisher), 3 µm CHIR99021 (Sigma-Aldrich), 1.25 mM N-acetylcysteine (Sigma-Aldrich), 10 mM Nicotinamide (Sigma-Aldrich), 10 nM recombinant gastrin (Sigma-Aldrich), 50 ng/ml EGF (Peprotech), 50 ng/ml FGF7 (Peprotech), 50 ng/ml FGF10 (Peprotech), 25 ng/ml HGF (Peprotech), 1 µM A83-01 (Tocris, Bristol, UK), 10 µM Rho Inhibitor γ-27632 (STEMCELL), and 1% growth factor-reduced matrigel. The same condition was used to culture PPTR tumor cells for two days to form iCCA spheroids.

4. **Hepatocyte 2D culture**

Freshly isolated HCs were subjected to 3D organoids culture as previously reported for five passages.[4] HC organoids were then dissociated and transferred to 60mm culture dishes precoated with 1% GFR matrigel and culture in 2D condition using the same organoid culture medium.

5. **iCCA-liver cocultures**

A 1:1 mixture of mouse HepatiCult™ Organoid Growth Medium (STEMCELL) and cholangiocyte organoid culture medium supplemented with 1% GFR matrigel was used in all cocultures.

(1) **WLC-T and HC-T spheroid cocultures (Figure 2B)**: WLC and HC spheroids were mixed with iCCA spheroids at a ratio of 15:1 in 96-well U-bottom microplates, and cultured for seven days using the same culture medium. The $ZsG^+$ tumor cell area was measured in Image J and plotted in GraphPad Prism 7.

(2) **iCCA-HC-MF 2.5D coculture (Figure 3 and Figure 4A)**: Freshly isolated *mTmG* HCs were mixed with primary mouse MFs at a ratio of HC:MF = 10:0 (HC-only), 8:2, 5:5, 2:8, or 0:10 (MF-only), seeded in 96-well flat-bottom microplates precoated with 1% GFR matrigel at

$2\times10^4$ total cells/well, and cultured in a 1:1 mixture of mouse HepatiCult™ Organoid Growth Medium (STEMCELL) and Stellate Cell Medium (ScienCell) for 24 hr. PPTR tumor spheroids were generated and added on the 2nd day at 2-5 spheroids/well. The cocultures were maintained up to 42 days and the medium was refreshed every 3-4 days. The ZsG fluorescence images were capture for the first four days, the diameters of the spheroids were measured in Image J and plotted in GraphPad Prism 7.

(3) **iCCA-HC-MF 2D coculture (Figure 4B)**: Two-chamber culture insert was placed in the µ-slide 8-well Grid-500 culture slides (ibidi, Gräfelfing, Germany) and the culture surface was precoated with 1% GFR matrigel at 37°C for 24 hr. PPTR tumor cells, MFs and 2D-cultured $TdT^+$ HCs were seeded in various combinations as indicated in Figure 4C. The total number of cells in each chamber was kept at $4.25\times10^4$ cells and the mixing ration of T:MF or HC:MF was 4:1. MFs were labeled with CellTracker™ Red CMTPX (Themo Fisher) in the conditions $TdT^+$ HCs were not used. The cocultures were maintained for 14 days and the culture medium *was* refreshed every *3*–4 *days.*

(4) **iCCA-HC 3D organoid coculture (Figure 5B)**: A 1:1 mixture of PPTR cells and HCs were seeded on top of 100% GFR matrigel in 48-well plate. The same spheroid coculture medium was used.

**iCCA Transplantation Models**

1. **iCCA orthotopic transplantation model**: PPTR-1 tumor cells were surgically injected into the liver of two-month-old male and female CD-1 nude mice. Briefly, a midline longitudinal abdominal incision was made to expose liver left lobe and PPTR-1 tumor cells were injected at $1\times10^5$/mouse in 4 µl cold growth factor-reduced (GFR) matrigel (Corning) using a Hamilton syringe and needle (Hamilton Company, Bonaduz, Switzerland). Survival curves and median survival of the orthotopic allograft models were determined by the Kaplan–Meier method in GraphPad Prism 7.

2. **Tail vein injection (TVI)-based iCCA lung metastasis model**: PPTR-1 tumor cells were injected into two-month-old male and female CD-1 nude mice at $1\times10^6$/mouse in 100 μl PBS via TVI.

**Cell Proliferation Assay**

PPTR cells were labeled by 5 μM CellTracker™ Red CMTPX dye (Invitrogen, Carlsbad, CA, USA) in Advanced DMEM/F12(Thermo Fisher) without FBS for 1 hour, then washed with Advanced DMEM/F12 (Thermo Fisher). 200 tumor cells/microwell and 100 PPTR + 100 MF cells/microwell in AW400 plates to aggregate to the spheroids. The culture medium was a mixture of 50% cholangiocyte organoid culture medium and 50% Stellate Cell Medium (ScienCell). HCs and MFs at a ratio of HC-only, 5:5, MF-only, and blank cultured in 96-well flat-bottom microplates. The culture medium was a mixture of 50% mouse HepatiCult™ Organoid Growth Medium (STEMCELL) and 50% Stellate Cell Medium (ScienCell). After 2 days, transferred around 2-3 spheroids/well into the 96-well plates. After 4 days culture, the cells have been collected and washed with 1x DPBS (Corning, Corning, NY, USA). Cells were resuspended with 100 μl and read the RFP using flow cytometry.

**Cytokine array assay**

Conditioned media (CM) were collected from cell cultures after four days. The levels of secreted cytokines in the CM were measured using Mouse Cytokine Antibody Array 3 (RayBiotech, GA, USA) according to the manufacturer's instructions. Briefly, 1 ml of undiluted CM were incubated with the Mouse Cytokine Antibody Array C3 membranes overnight at 4°C. After washing, the membranes were incubated with the Biotinylated Antibody Cocktail and subsequently with HRP-Streptavidin, 2 hours at room temperature for each. After washing, the cytokine signals were visualized via the LI-COR chemiluminescence imaging system. The signals were quantified using Image Studio Lite Ver5.2 (LI-COR) and normalized to the positive controls on the membrane.

**Cell Migration Assay**

Cell migration capacity were measured using polycarbonate membrane inserts (8 μm pore size, Cell BioLabs, USA) in 24-well plates. In brief, 2 different PPTR cell lines (PPTR1 and PPTR2) were seeded in same density in 60mm dish and treated with 5 μM Vcam1$^{Ab}$ (Bioxcell, Lebanon, NH, USA) and IgG (R&D System, Minneapolis, MN, USA) respectively for 4 days. Then, PPTR-1 and PPTR-2 cell suspension were added to the upper chamber (3 ×$10^4$ cells in 300 μl blank medium with 5 μM of Vcam1 antibody or IgG respectively), and 500 μl of media with 10% FBS was added to the lower chamber. After incubation at 37 °C under 5% $CO_2$ for 24 h, the cells on the upper surface were wiped with cotton swabs, while the cells that migrated through the filter pores were stained with cell stain solution for 10 min. The stained cells were observed and photographed under an inverted microscope. Cells were counted in five randomly selected fields of view using Image J.

**Image-based quantification**

1. **Tumor cell proliferation (Figure 1D):** 200x Ki67 IHC images were captured from three independent tumor areas for each group. The number of nuclei and Ki67$^+$ cells were manually counted and plotted in GraphPad Prism 7. Student *t* test was performed for comparison.
2. **Tumor cell expansion in spheroid coculture (Figure 2C):** The ZsG$^+$ tumor cell area was measured in Image J and normalized to Day 0 area as fold-change, then plotted in GraphPad Prism 7. Student *t* test was performed for comparison.
3. **Tumor cell expansion in the two-chamber coculture (Figure 4C):** ZsG/bright-fiend images of the cocultures from Day 0 and 14 were aligned in Photoshop using the 500-μm engraved grid on the culture slides. The movement distance of the tumor border was measured from three independent sets of experiments in Image J and plotted in GraphPad Prism 7. Student *t* test was performed for comparison.
4. **Loss of hepatocyte area in two-chamber coculture (Figure 5E):** Area occupied by TdT$^+$ hepatocytes within the original seeding space on Day 0, 7, 14, and 21 was measured from three

independent sets of experiments in Image J, normalized to Day 0, and then plotted in GraphPad Prism 7. Student *t* test was performed for comparison.

5. **Lung metastasis area measurement (Figure 6F, G):** Whole lung ZsG fluorescence images were captured by AxioScan Z.1 Whole Slide Scanner (Zeiss, Oberkochen, Germany). The number of lung metastases and the area of individual metastasis were measured in Image J and plotted in GraphPad Prism 7. Student *t* test was performed for comparison. Three lungs each from the IgG and $Vcam1^{Ab}$ group were compared with the similar results.
